## Supplementary figure legends for "Huntingtin interactome reveals huntingtin role in regulation of double strand break DNA damage response (DSB/DDR), chromatin remodeling and RNA processing pathways"

### Supplementary figures legends

**Figure S1. Pan-nuclear γH2A.X staining of apoptotic cells, not associated with DSB.** Examples of apoptotic cells (~5-10%) not included in analysis. IF staining with γ-H2A.X specific antibody. Nuclei were visualized with DAPI

**Figure S2. Detection and relative quantitation of phosphorylation on HTT from ISPNs using Parallel Reaction Monitoring (PRM).** Peak Areas for precursors and products (fragment ions) for non-treated (NT) and bleomycin-treated 33CAG and HD 180CAG ISPNs are shown on the left. Unmodified peptide from HTT were used for normalization to account for the amount of HTT in samples. Transitions (m/z ratio of a peptide and its corresponding product ion m/z) are shown on the right for selected phosphorylated tryptic peptides.

**Figure S3. HTT /TCERG1 co-localization and interaction in ISPNs** A, Representative images of ISPNs treated with bleomycin (10μg/ml, 2 h) co-stained with TCERG1G1 and HTT (MCA2050) antibodies. B, Representative co-IP experiments from control (33) and HD (180) ISPNs treated with bleomycin (10μg/ml, 2h) and untreated (NT). Total cell lysates were prepared and HTT complexes were immuno-precipitated using antibodies to total HTT (MCA2050). TCERG1 and HTT proteins were detected in the IPs. IgG negative control IPs are shown at the bottom panel. The inputs for HTT, DNA-PKcs and beta-tubulin are shown (right panels). C, Proximity Ligation Assay (PLA) in normal (33CAG) and HD (180CAG) ISPNs treated with bleomycin (10μg/ml, 2 h) and untreated using TCERG1 and HTT antibodies. Images shown for 180 ISPNs were taken at higher intensity for illustrative purpose, while quantitation was done with the same settings as for 33 ISPNs. Graphs (D) show quantification of PLA signals using MetaXpress software (Molecular Devices). The data is presented as mean $\pm$SEM of the number of nuclear PLA sites per cell, sum intensity of nuclear PLA sites per cell and average nuclear PLA site intensity relative to technical negative control within each experiment. 30-50 cells were analyzed for each condition. One-way ANOVA with Pairwise Multiple Comparison Procedures (Holm-Sidak method): *$p<0.001$, n=3; **$p=0.01$, n=3. T-test with equal variances: ^$p<0.001$, n=3; ^^$p=0.005$, n=3; $^{\#}p=0.003$, n=3; ## $p=0.016$, n=3

**Figure S4. Nuclear HTT is depleted in HD ISPNs.** Representative images of ISPNs treated with bleomycin (10μg/ml, 2 h) and untreated co-stained with HTT (MCA2050) and Golgi marker GM130 antibodies. HD cells with depleted nuclear HTT are indicated by arrows.

**Figure S5. Optimization of APEX2-mediated proximity biotin labeling for HTT interactions**. Representative experiment is shown. HEK293 cells were transfected with APEX2-Htt-22Q plasmid. Biotin-labeling was performed as described in the Methods Section. No biotin (lane 2) was added for negative control. Biotinylated proteins were enriched by streptavidin pulldown, eluted by incubation in 2× SDS protein loading buffer (Biorad) supplemented with 2.5 mM biotin and 20 mM dithiolthreitol (DTT) for 10 min at 95 degrees, and used for western blot. Biotinylated proteins were detected using streptavidin-HRP (Cell Signaling Technology) in the inputs and elutions (but not in negative controls). Fewer biotinylated proteins in flow-through indicates of efficient binding to streptavidin resin.
