## Supplementary figures and images for "Huntingtin interactome reveals huntingtin role in regulation of double strand break DNA damage response (DSB/DDR), chromatin remodeling and RNA processing pathways"

### Fig. S1

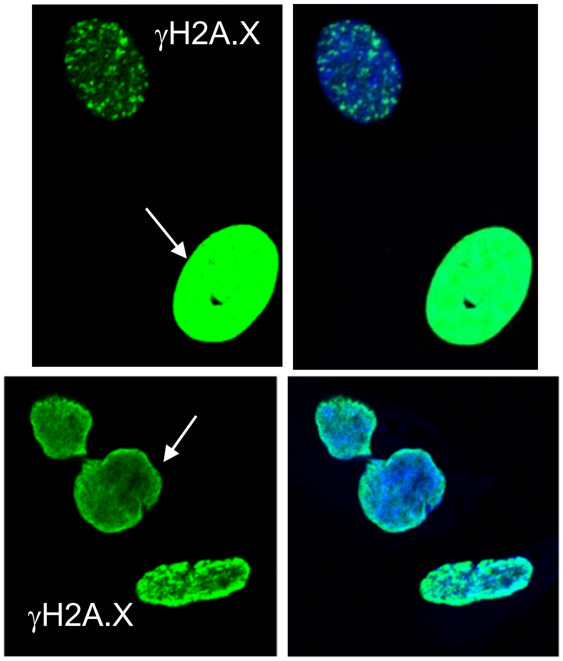


Figure S1

### Fig. S2

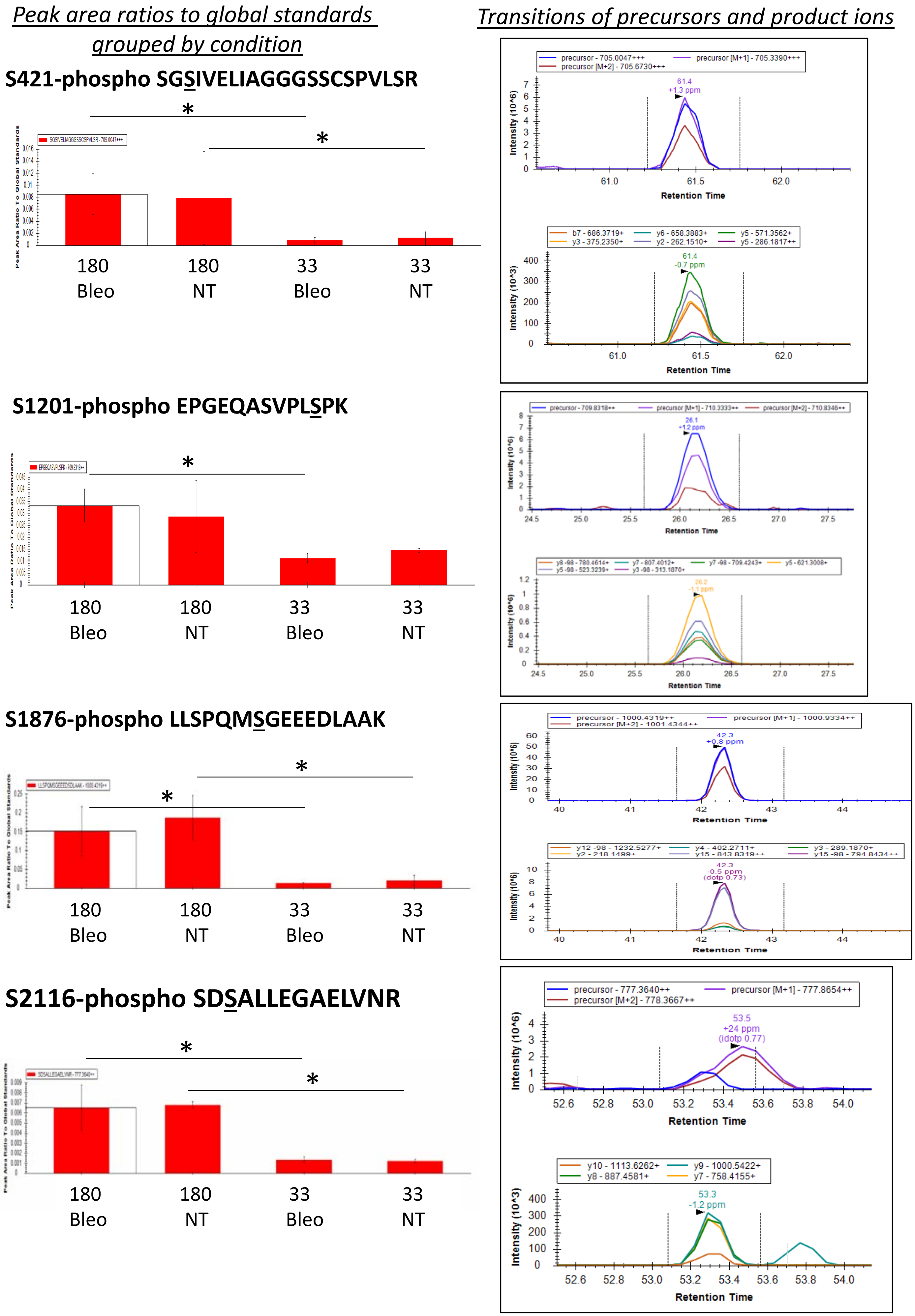

### Fig. S3

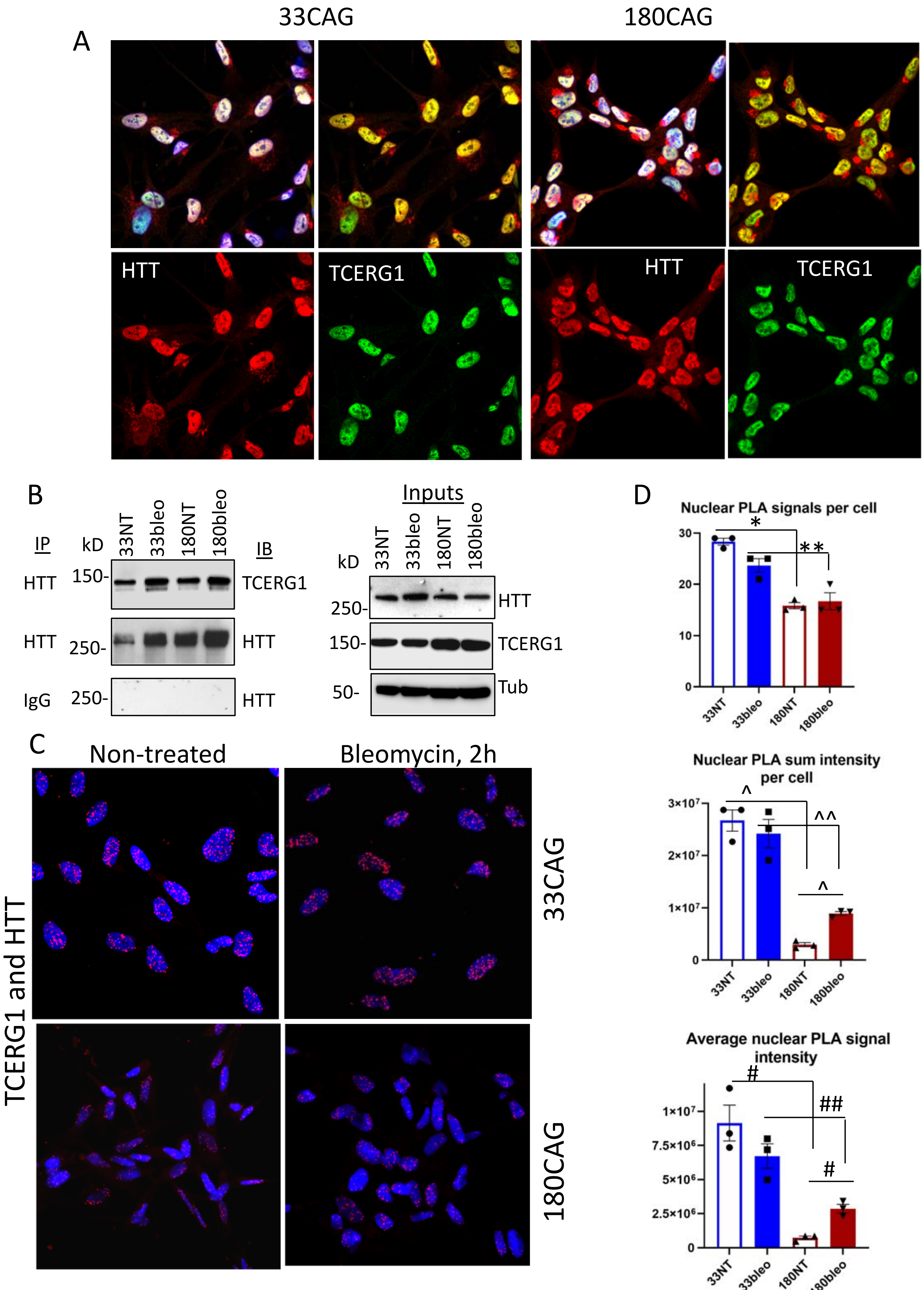

### Fig. S4

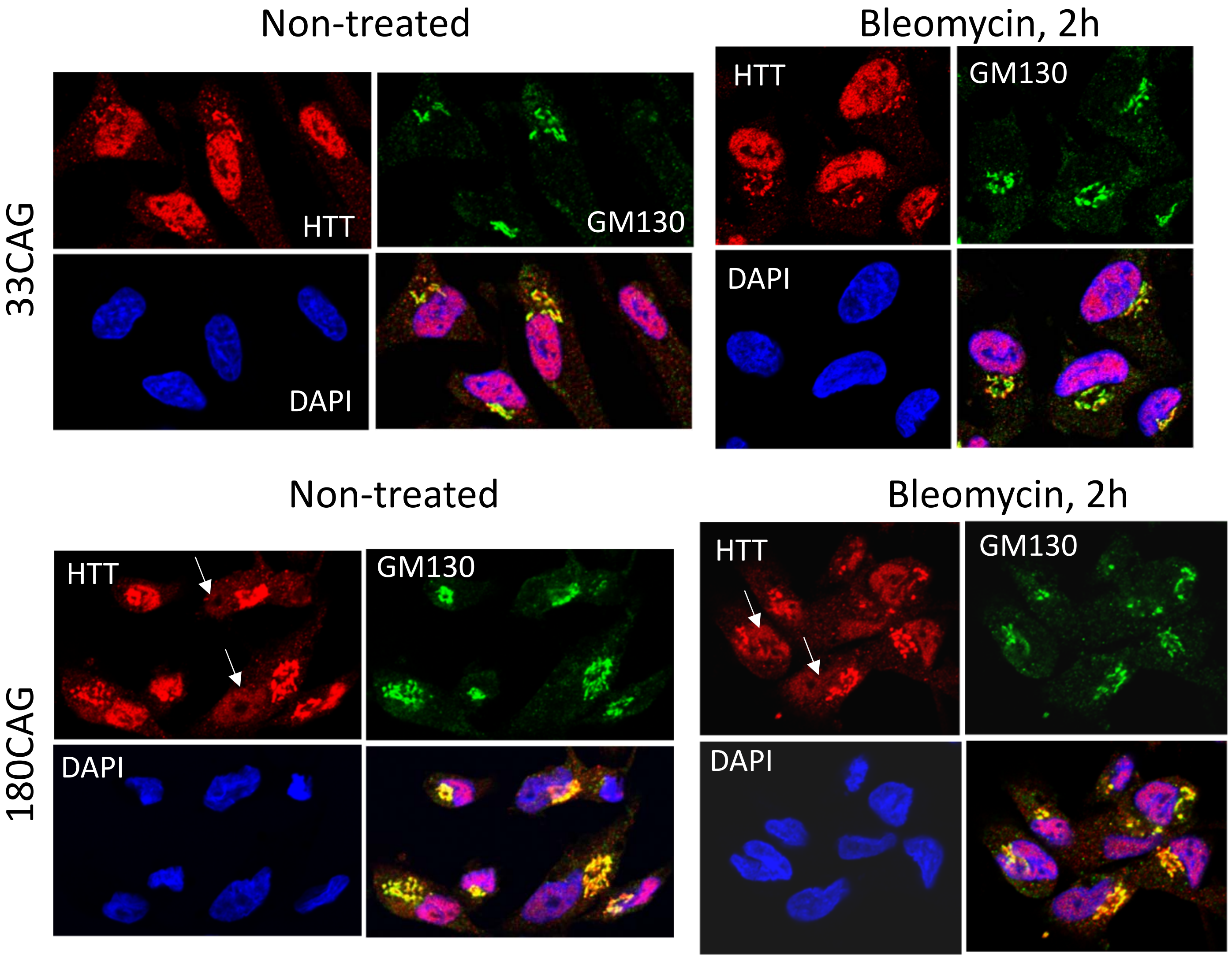

### Fig. S5

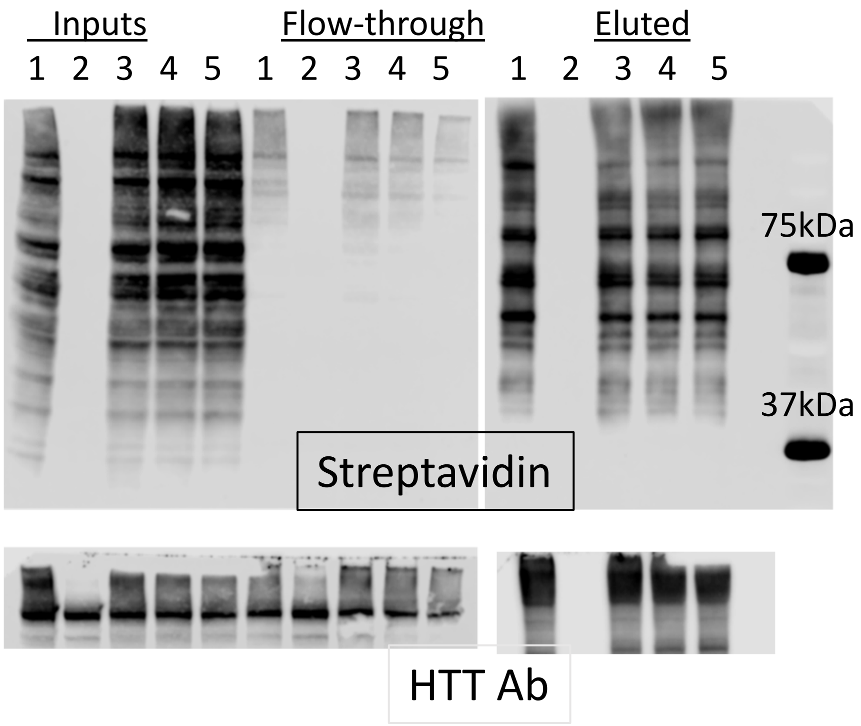
